# Nested Connections: Local Phage and Broad Plasmid Sharing in the Honey Bee Mobilome

**DOI:** 10.1101/2025.10.31.685865

**Authors:** Delaney L. Miller, Lílian Caesar, Carrie Ganote, Sergio López-Madrigal, Danny W. Rice, Chris R. P. Robinson, Alyssa M. Seawright, Jillian P. Lewis, Irene L. G. Newton

**Affiliations:** University of Wisconsin Madison; Indiana University Bloomington; University of Connecticut

## Abstract

Honey bees rely on bacterial symbionts for their nutritional needs and for protection against invading pathogens. Genetic diversity among strains within the colony has the potential to impact symbiont function and subsequently the benefits that honey bees receive from them. Mobilized genes vectored by mobile genetic elements (MGEs) like phages, transposons, conjugative elements and plasmids, are known to rapidly alter bacterial phenotypes. We identified phages and plasmids in genomes of the symbiont *Bombella apis* between two colonies, with the goal of understanding which MGEs contribute to strain diversification as well as MGE distribution across colonies and between microbial species. Interestingly, we found some *B. apis* strains carry plasmids while all harbor a diversity of integrated phages, with only one phage clusters conserved across all. Identified *B. apis* phages are not found outside of the *Bombella* and *Saccaribacter* species, suggesting some host specificity for these MGEs. Of the five plasmids discovered, two appear to be phage-plasmids with high similarity to phages found in previously sequenced *B. apis* genomes. Interestingly, three plasmids in *B. apis* shared significant average nucleotide identity with known plasmids from acetic acid bacteria isolated from flowers, plants, and fermented foods. This result suggests that *B. apis* has acquired MGEs, either vertically or horizontally, from plant- and fermented-food associated AABs. Overall, our findings suggest that MGE content varies between colonies and has the potential to shape genetic and phenotypic variation between strains.

## Introduction

Bacteria are ubiquitous among eukaryotic hosts and their role in shaping host health is a tenet of modern biology, and yet we still understand only a fraction of their genetic and functional diversity [1, 2]. Pangenome analyses of bacteria, which aim to capture the breadth of genes that vary between strains within a species, show that symbionts with free-living life stages have highly variable or “open” pangenomes, meaning they are more variable due to gene gains and losses [3]. This fluidity can have consequences for host health. Strain-level variation among symbionts is known to affect disease outcomes, nutrition availability, and host development [4–6]. It is therefore important to use population-level approaches to identify gene gains and losses that may impact symbiont function.

Among bacteria, mobile genetic elements (MGEs) such as plasmids, insertion sequences, integrative conjugative elements (ICEs), and phages facilitate rapid gene gain and loss [7]. Among symbionts, the most well-studied examples of mobilized traits can be found in plant and insect symbionts. A classic example is rhizobial symbionts, which cannot fix nitrogen for their legume hosts without the nitrogenases encoded on MGEs (plasmids or ICEs depending on the species) [8–10]. In insect symbionts, the gene clusters that encode for antimicrobials, which defend their host from disease, are often mobile, horizontally transmitting between species [4, 11]. Another striking example of host traits mobilized by bacterial MGEs is from the symbiont *Wolbachia pipientis* and its prophage WO which carries a two-gene module responsible for cytoplasmic incompatibility, a mechanism for reproductive manipulation [12]. Clearly, symbionts and their myriad MGEs have the potential to radically change their host’s health and ecology, but MGEs remain understudied, both due to the challenges recovering them from genome assemblies and the difficulty classifying them, especially in non-model bacterial hosts [13–15].

Honey bees are agriculturally important pollinators and their microbiomes impact bee health and productivity [16–18]. Within the adult honey bee microbiome, strain variation impacts important traits, from pathways that detoxify sugars in the bee’s diet to the T6SS, which underpins interbacterial competition [19, 20]. These strains often carry many different mobile genetic elements (MGEs) [21–24] such as phages and plasmids, which are often shared between bacterial species [24] and carry cargo genes with metabolic functions predicted to benefit honey bees [25]. In other colony environments, such as the shared food reserves or the guts of larvae or queens[26], it is less understood what MGEs are present and how they impact symbiont function.

Here, we survey MGEs found in *B. apis*, a symbiont that colonizes multiple colony environments, from larvae and queens to the colony’s food reserves. Since *B. apis* is exposed to different microbial communities in these environments, we hypothesized that strains from different environments would also carry different MGEs. In particular, we expect that genetic exchange between honey bee symbionts and environmental microbes is most likely to occur at the interface between host digestive tracts and foraged resources: the food reserves (Figure 1B). Nectar and pollen are collected from nearby flowers by foragers, stored in cells at the periphery of the hive (Figure 1A), and fed to the rest of the colony. At the center of the colony, where the queen lays her eggs, nurse bees feed (Figure 1C) the nectar to the queen and larvae by regurgitating from their foregut, also called the crop [27]. In addition to nectar and pollen, queens and young larvae are fed royal jelly, a secretion of transformed pollen from the nurses’ hypopharyngeal gland (Figure 1D & E) [28]. This flow of resources from the food reserves at the periphery of the hive, to nurses and subsequently the larvae and queen is thought to reduce exposure of larvae and queens to environmental microbes. As nectar and pollen flow to the center of the colony, they are transformed in ways that likely exclude environmental microbes. In the food reserves, nectar stored long-term is dehydrated into honey, and pollen is fermented into bee bread [27, 29]. These changes in osmotic pressure and acidity create environments inhospitable to many microbes. Any microbe capable of surviving in the food reserves may be picked up by a nurse bee and fed to the queen or larvae, along with a serving of royal jelly, which is highly toxic to most bacteria due to antimicrobial peptides, low pH, and viscosity [30, 31]. This gauntlet of transmission to increasingly inhospitable colony environments means that most transient, environmental microbes do not make it past the food reserves [32–34]. Therefore, it is likely that honey bee symbionts will have the best chance to acquire MGEs when in the food stores, where foragers deposit floral resources with foreign microbes. Among the microbes that are capable of surviving in nectar, and the fermented food that honey bees and many other pollinators eat, are acetic acid bacteria (AAB) and lactic acid bacteria (LAB) [35, 36].

**Figure 1:**
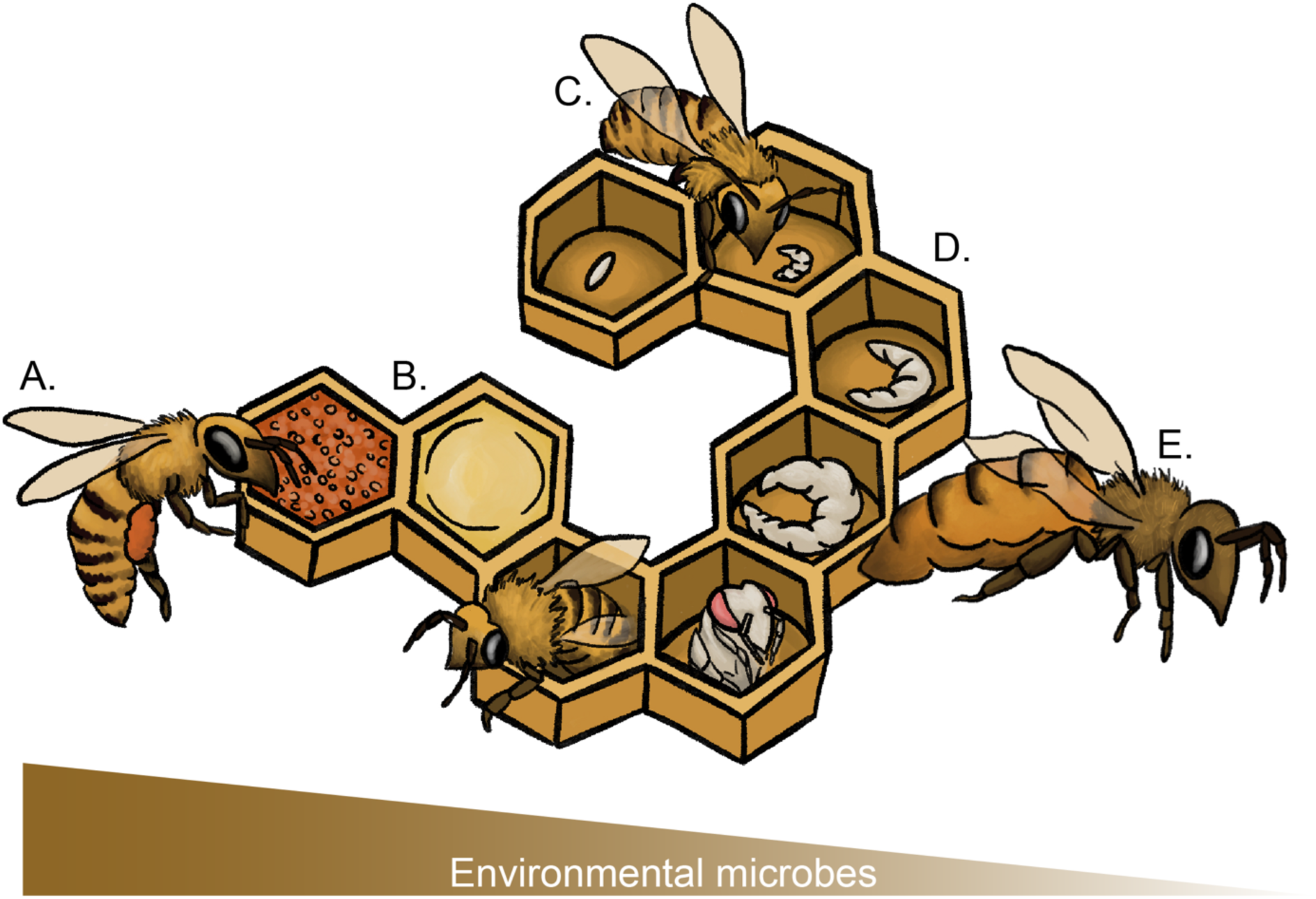
Distribution of *Bombella apis* across colony environments: (A) Foragers visit flowers and transport nectar (via crop) and pollen (via pollen baskets) back to the colony. The influx of nectar and pollen (B) brings in exogenous microbes: nectar and pollen associates, as well as microbes deposited by other visiting pollinators. These microbes transiently persist in the nectar and pollen, which are incorporated into the diet fed to larvae (D) by nurse bees (C) and to (E) queens by her attendants. Microbial diversity drops in the diet, likely due to the addition of antimicrobial royal jelly (synthesized by nurses and attendants).

This combination of factors, described above, that eliminate transient microbes, may explain the low species diversity of the nectar, larval and queen microbiomes. All three microbial communities are dominated by AAB and LAB, including *Bombella apis* and *Apilactobacillus kunkeeii* [33, 34, 37] in the nectar and larvae, and *B. apis* and a consortium of LAB in queens [26, 38]. Both *B. apis* and *A. kunkeeii* have been shown to have defensive properties against pathogenic fungi and bacteria respectively [17, 39]. *B. apis* is also a nutritional symbiont associated with both queens and larvae, and survives in the presence of high concentrations of royal jelly, which are toxic to *A. kunkeeii* and other microbiome members [18]. *B. apis* is unique in its distribution within the colony, as it is predominantly found in the queen digestive tract, larval gut, nurse hypopharyngeal glands, nurse crops, and foraged resources. *B. apis* therefore exists in niches that have either high or low exposure to environmental microbes. In this study we isolated new *Bombella* strains across colony environments – from nectar to crops to queens and larvae – with the aim to understand how MGEs have shaped their genetic diversity.

We sequenced the isolates’ genomes, generated complete and closed assemblies, and used those genomes to ask the following questions about plasmids and phages we identified: (1) What MGEs are shared between *B. apis* strains? (2) What cargo genes do they encode that might shape *B. apis*’ phenotypic diversity? (3) Lastly, which bacterial species, both inside and outside the colony, share similar MGEs? Ultimately, we identified many phage clusters unique to our apiary, and one phage cluster conserved across most *B. apis* genomes. Plasmids were rare, found in only five genomes and not shared between *B. apis* strains. Despite sharing low genetic similarity, the plasmids all carried toxin-antitoxin systems, and two plasmids (pJPL21 and pAML2) encode phages with high similarity to those found in other *B. apis* genomes. Phage-plasmids, pJPL21 and pAML2, have CDSs similar to other *Bombella* and *Saccharibacter* genomes, whereas plasmids (pJPL24_01, pJPL24_02, and pSME1) had backbones and cargo genes more similar to plasmids from more distantly-related AAB. In short, phages vary colony to colony and have the potential to shape genetic and phenotypic variation between *Bombella* strains. Plasmids, on the other hand, are rare among *B. apis* strains and their similarity to other AAB isolated from plants, other insects, and fermented foods, suggests they may have been acquired from AAB from outside the colony.

## Materials and methods

### Strain collection and identification

Late instar larvae, nurse crops, and queen guts from two bee colonies in the Indiana University research apiary were suspended in 1X PBS, plated on MRS media and incubated for 48 hours at 34° C. Isolated bacteria were cryoarchived, extracted with Qiagen’s MagAttract HMW DNA kit, PCR amplified (using 27F, (AGAGTTTGATCMTGGCTCAG) and 1492R (TACGGYTACCTTGTTACGACTT) primers) and Sanger sequenced at QuintaraBio. Top NCBI BLASTn [40] hits (>99% sequence identity) was used to identify species.

### gDNA extraction and long-read sequencing

Nineteen new isolates and one previously identified isolate were chosen for long-read, whole genome sequencing. gDNA extractions followed Quiagen’s MagAttract HMW DNA’s gram-negative protocol. Quality control of extractions was determined with a Qubit fluorometer and Nanodrop spectrophotometer. gDNA was sheared to approximately 13kb fragments with a Megaruptor 3 sonicator and libraries were prepared using the SMRTBell Express Template Prep Kit 2.0. One SMRTcell 8M was used to sequence libraries on the PacBio Sequel lle on CCS sequencing mode and a 30hs movie time. Circular Consensus Sequence (CCS) analysis was performed on the instrument using SMRTLink V11 (parameters: ccs –min passes 3 –min rq 0.99) to obtain HiFi reads.

### Sequencing QC, assembly, and annotation

HiFi reads were checked for quality using longQC (parameter: -x pb-sequence) [41]. Any samples with more than one GC peak were removed. Reads were assembled with Canu (parameters: genomeSize=2M -pacbio-hifi) [42] three times to produce three distinct assemblies (since Canu is non-deterministic) which were used to generate a consensus assembly with Trycycler [43]. Default parameters were used unless otherwise stated. Contig clusters were removed if found in only one assembly. During reconciling, clusters with mismatched lengths were excluded. Consensus assemblies were used for all downstream analysis. Annotation was done with NCBI’s PGAP software version 2022-08-11.build 627; this annotation was used in later analysis unless otherwise stated [44]. For already published genomes, the NCBI entries annotated with PGAP were also used so that annotations would be consistent across genomes for ortholog analysis. This reduces false positive gains or losses due to different gene calling software.

### Core ortholog phylogeny

Proteins were clustered into ortholog groups by requiring all proteins in a cluster to be reciprocal best BLASTP [40] hits with all other proteins in the cluster (i.e. complete linkage clustering of reciprocal best hits). In addition, if two or more proteins within a genome were more similar to each other than they were to any protein in any other genome (likely duplications within the lineage leading to that genome), all such proteins were placed into the same ortholog group. Protein alignments of ortholog group members were calculated with LINSI [45]. The corresponding codon alignments were derived from these protein alignments and concatenated into a 1.8 MB alignment. Before concatenation, a filter was used to exclude some ortholog groups based on the following criteria: 1) The group had to include sequences from 20 or more of the genomes, and 2) The median all vs all pairwise amino acid identity had to be 60% or higher. A pairwise amino acid identity was defined as (number of identical amino acids) / max(seq1_len, seq2_len) * 100. This last criterion was to reduce misalignments of divergent orthologs, groups with large disparities in group member length, and ortholog groups containing non orthologous genes. All available nucleotide evolution models were tested using IQTREE [46]. The best model found based on the Bayesian information criterion was the GTR model with invariant sites and a gamma rate distribution using 30 categories. The final tree was calculated with this model along with 1000 bootstrap replicates in IQTREE.

### Gene gain/loss analysis

Gene gains and losses were inferred for each ortholog group relative to the species tree based on maximum parsimony with the cost of a gain set to 2.001 times the cost of a loss. The gainLoss program used by the GLOOME web server [47] was run locally for this analysis.

### COG category assignment

A representative sequence (based on median length) from each ortholog group was BLAST-ed [40] against COG member proteins (3,213,025 sequences) from the 2020 COG database [48]. Only COG member proteins in “COG membership classes” 0 and 1 were considered, as these member proteins cover most of the COG. For a *Bombella* ortholog to be assigned to a COG category it had to overlap ≥ 60% of the COG members range that aligned to its COG, meaning that in most cases the *Bombella* protein would align with ≥ 60% of the COG itself, give the “COG memberships class” constraint above. The highest BLASTP bit scores were considered first. If lower scoring matches to different COGs existed, the *Bombella* protein could be assigned to multiple COG categories if the above constraints were met and the *Bombella* region matching such a COG did not overlap with regions already assigned to a COG by more than 60%. Of the 5924 *Bombella* orthologs that were assigned to some COG only 17 were assigned to 2 COGs and none to more than 2.

### Curation of plasmid annotation

Plasmid annotation was manually refined as follows: the sequence similarity of all loci and intergenic sequences (IGSs) were searched in NCBI’s non-redundant protein sequences (nr) database through Blastx using default parameters [40]. Automatic annotation artifacts (e.g. split loci, unannotated loci) were corrected when detected. Endogenous replication initiator proteins (i.e. either repA, repB, or repC, depending on the plasmid) were set as the first nucleotide in the assembly. Final GenBank files were fed to the Proksee web server [49] in order to generate plasmids’ graphical maps. Categorization of coding sequences by COG functional category was completed with eggNOG [50].

### Plasmids visualization in electrophoresis

All *Bombella* strains were grown overnight in 4 mL MRS (34° C, 250 rpm). Liquid cultures were split downstream for either total DNA extraction or plasmid DNA extraction with Qiagen kits (DNeasy or MiniPrep). Approximately 150 ng DNA resulting from each extraction protocol were analyzed in a 0.5% agarose gel electrophoresis (80 Volts, 100 min).

### Plasmid and phage coding sequence similarity analysis

Plasmid and phage coding sequences were compared to all publicly available genomes on NCBI using cblaster [51], setting a minimum query coverage and percent identity to 50%. To increase the stringency of our search we further modified the parameters. Namely we required that at least 25% of plasmid coding sequences were present in a neighborhood less than or equal to 1.5 * length of the query plasmid or phage. This threshold was determined for the following reasons: (1) plasmid sequences misassembled into bacterial genomes will be linear sequences, and therefore genes on opposite ends of the contig may actually be neighbors; (2) plasmids and phages expand and contract in length frequently as they acquire or lose new DNA. By allowing for a 50% increase in size, we can capture similar plasmids or phages that have expanded in length. The amino acid similarity and synteny of plasmids with similarity to *B. apis* plasmids were visualized with gene arrow maps using clinker [52].

### Plasmid nucleotide sequence similarity analysis

Sourmash [53] was used to calculate pairwise Jaccard similarity between query plasmids and curated databases of plasmid sequences. Plasmid databases were compiled from one of three sources: (1) PLSDB sequences isolated from honey bees, (2) the top 100 tBLASTn [40] hits to each plasmid’s replication initiation protein, and (3) cblaster hits from verified plasmid sequences. In sourmash, k-mers of 31 bp were used, as recommended for the genetic relatedness between our input sequences.

### Identification, annotation, and validation of prophages

All *B. apis* genomes, including genomes recovered from NCBI, were screened for the presence of prophages with VIBRANT [54]. Recovered prophages were annotated using DRAMv [54] Annotations were then queried for the presence of canonical genes for prophage excision [55] Prophages without any canonical gene for prophage excision were classified as degenerate while prophages with canonical excision genes and at least one structural gene were classified as complete. Only prophages with sufficient genome quality (Medium or higher) were considered.[56] Recovered prophages were then clustered into genus and species-level OTUs with dRep [57] dRep dereplicate -l 2000 --ignoreGenomeQuality -pa 0.8 -sa 0.95 -nc 0.85 -comW 0 -conW 0 -strW 0 -N50W 0 -sizeW 1 -centW 0). These parameters are informed by observations of known phage biological and ecological diversity [58]. Prophage-encoded auxiliary metabolic genes (AMGs) were inspected by comparing the output of VIBRANT and DRAM-v.

## Results

### Closely related strains co-occur within the same colony

To compare the genotypic diversity of *B. apis* strains between bee colonies and within colony environments, bacteria were isolated from either nectar (Fig. 1B), nurse crops (Fig. 1C), larvae (Fig. 1D), or queen guts (Fig. 1E), between two bee colonies and identified as *B. apis* by 16S sequencing. The resulting nineteen new isolates (Table 1) - as well as SME1, a nectar isolate collected four years prior for which we only had short-reads [59] - were long-read sequenced on PacBio’s Sequel II platform with an average of ∼300x coverage and assembled with a combination of Canu and Trycyler [42, 43] (Table 1). To determine the phylogenetic placement of our new genomes, an ortholog tree was inferred (Fig. 2A-B), using all publicly available *Bombella* genomes and other closely related insect and flower-associated AAB as outgroups. Of the nineteen new *Bombella* strains, two larval strains were placed within the *Bombella favorum* clade, a species previously identified in honey bee nectar provisions [60] (Fig. 2A). *B. apis* genomes were GC rich (∼59%), as expected, with lengths around 2 Mb (Table 1). Genome length across the dataset varied by up to 100,000 bp, ranging from 1.98 to 2.2 Mb in total, hinting at the gain or loss of large regions.

**Figure 2:**
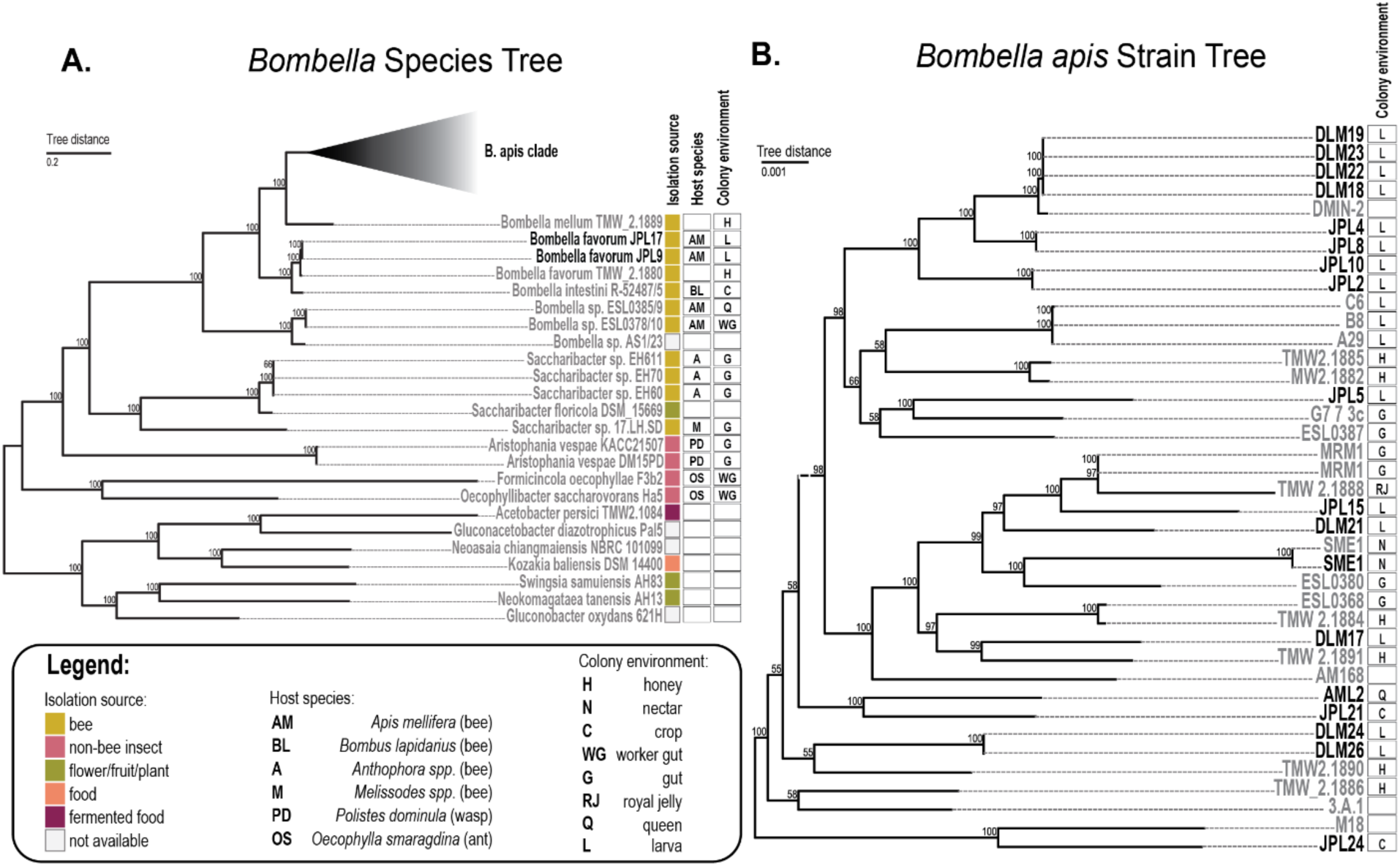
*Bombella* strains isolated from the same colony and/or environment are phylogenetically distant: A. Core ortholog phylogeny of all sequenced *Bombella* and (B) *Bombella apis* genomes and closely-related acetic acid bacteria from insects, plants, and fermented foods. Phylogeny was inferred from 2,711 orthologs (see Methods) using IQ-tree. New genomes generated from this study are highlighted in black. Isolation source is denoted by colored boxes, and, when applicable, the host species and colony environment are labeled with acronyms.

**Table 1:**
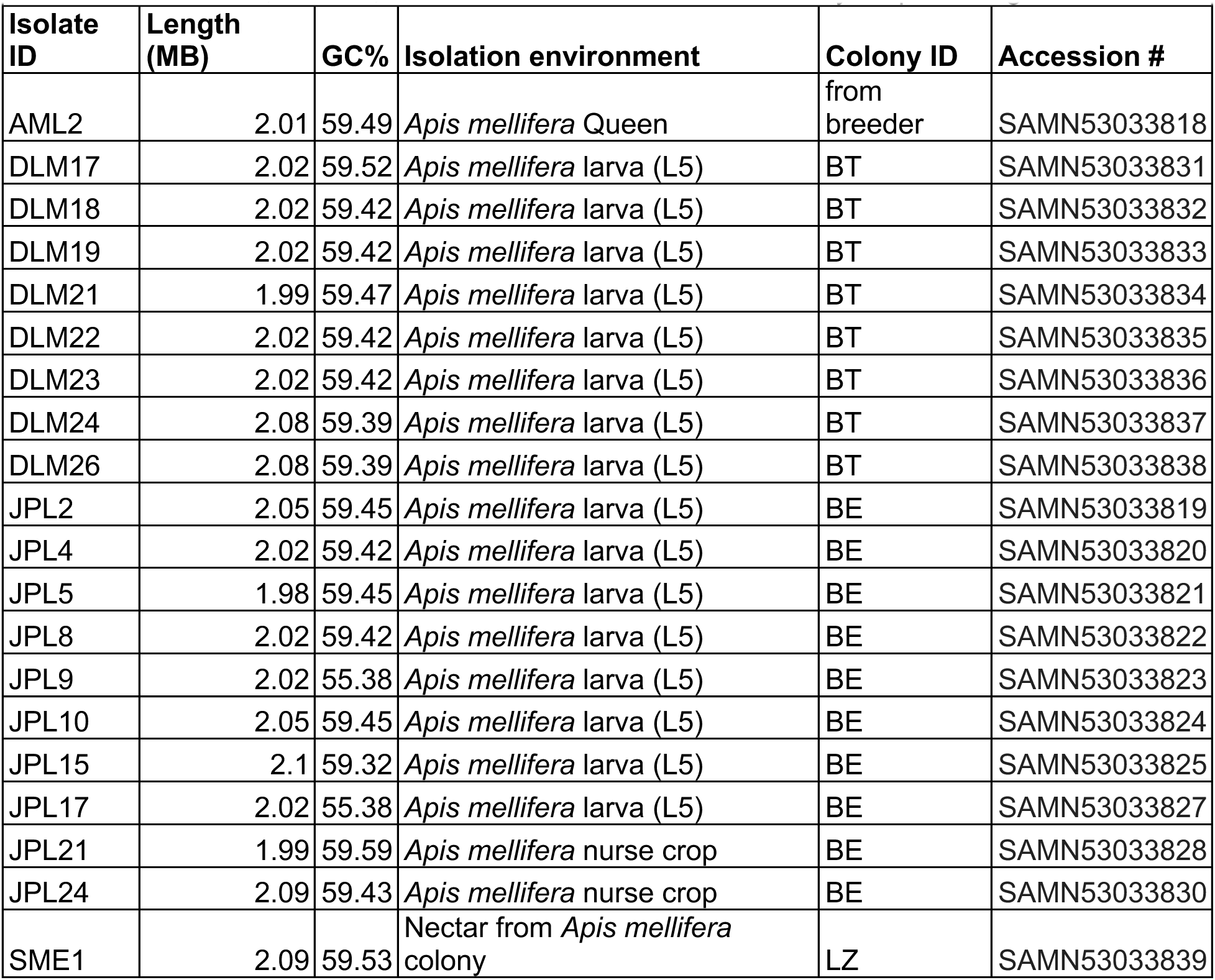
Genome ID, metrics, and isolation environments for newly-sequenced genomes.

Some strains isolated from the same colony (colony 1: JPL; colony 2: DLM) were closely related and formed monophyletic clades (i.e. DLM19-18) (Fig. 2B). Other strains were more closely related to isolates from Europe (i.e. ESL0387 from Switzerland and JPL5; TMW2.1891 from Germany and DLM17). In short, strains sampled from our apiary were diverse, and spanned the *B. apis* phylogeny. It is worth noting that *B. apis* strains are all highly related - regardless of isolation source. The average nucleotide identity (ANI) across all strains is ∼99%.

### Recent gene gains and losses are driven by mobile genetic elements and defense islands

To better understand the diversification of strains within the same colony and/or apiary, we identified genes gained or lost within the *B. apis* clade using GLOOME [47]. Specifically, we were interested in the most recent gene gains - genes that are unique to each new strain we sequenced. To better visualize the broad pattern of what functions are newly gained and lost, we classified gained/lost genes by their COG (Cluster of Orthologous Groups of proteins) category (24) (Fig. 3; Fig. S1). Across the *B. apis* tree, 24.3% of gene gains were classified by COG as mobile elements. Gene gains unique to our strains were largely involved in replication and repair (L), defense mechanisms (V), and mobile elements (X) (Fig.3A). Gene gains due to mobile elements were largely driven by phages (Fig. 3C, suggesting that phages are contributing to genetic diversity among strains of the same colony.

**Figure 3:**
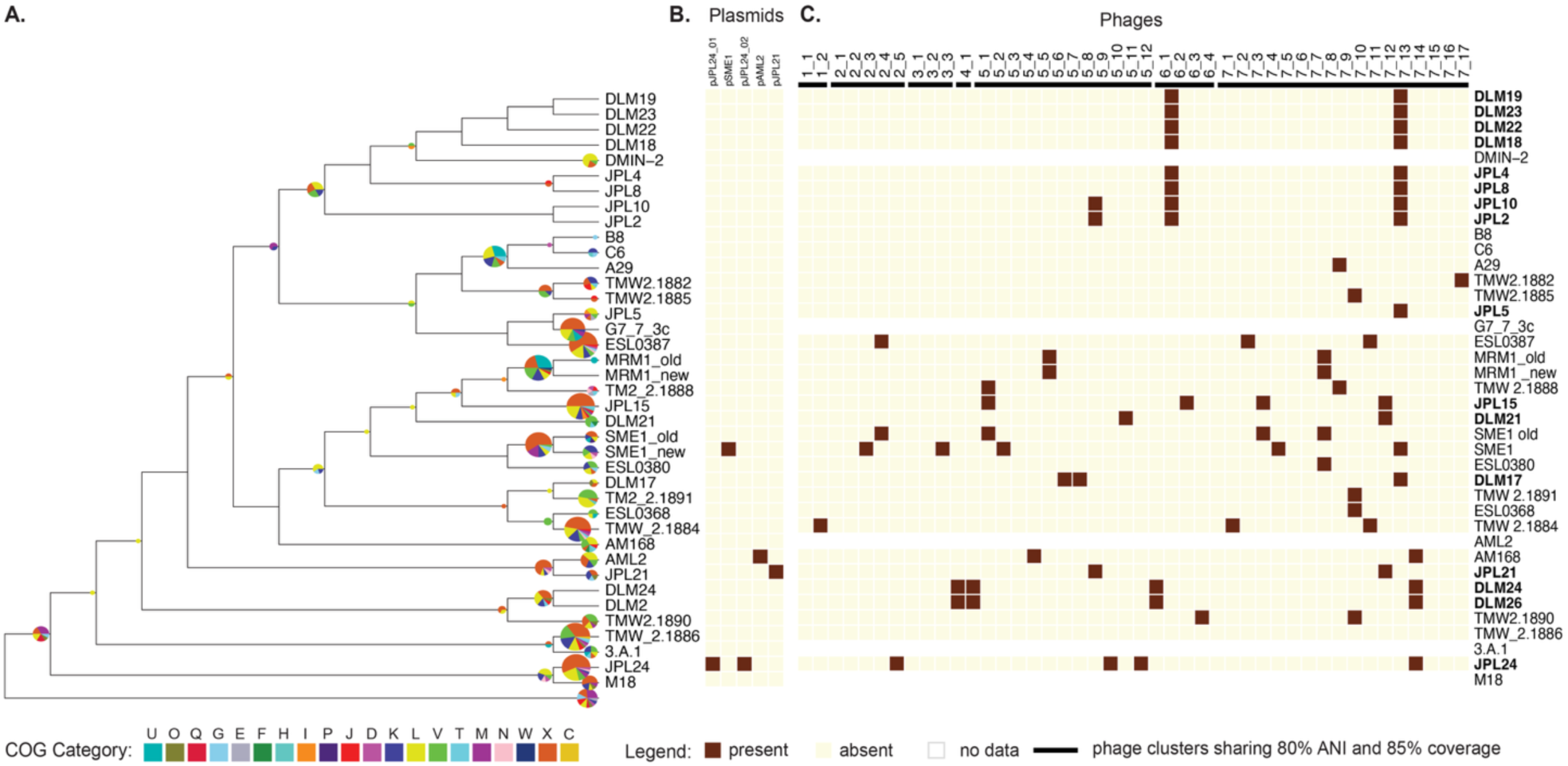
Strain-specific gains are biased towards mobile elements and defense islands: (A) A. COG categories of genes gained within the *B. apis* clade. Gene gains and losses were defined with GLOOME and categorized by their COG family. Pie charts denote the COG categories of genes gained at that node or leaf. Pie chart colors represent COG categories (U: intracellular trafficking, secretion and vesicular transport; O: posttranslational modification, protein turnover and chaperones; Q: secondary metabolite biosynthesis, transport, and catabolism; G: carbohydrate transport and etabolism; E: amino acid transport and metabolism; F: nucleotide transport and metabolism; H: coenzyme transport and metabolism; I: lipid transport and metabolism; P: inorganic ion transport and etabolism; J: translation, ribosomal structure and biogenesis; D: cell cycle control, cell division, and chromosome partitioning; K: transcription; L: replication, recombination, and repair; V: defense mechanisms; T: signal transduction mechanisms; M: cell wall/membrane/envelope biogenesis; N: cell motility; W: extracellular structures; X: mobilome, prophage, and transposons; C: energy production and conversion) and the diameter of each pie represents the number of genes gained. (B) Plasmid presence. Presence of at least one plasmid identified during genome assembly is denoted in filled squares. (C) Presence of phage within each genome. Phage presence and absence was annotated with Vibrant. Phages with incomplete integrase and structural capsid genes are considered degenerate. VIRDIC was used to bin phage sequences at the genus level, identifying 14 distinct genera present in this clade.

### Identification of phages within *B. apis* genomes sampled from different colonies

To identify groups of related phages – termed clusters – shared between *Bombella* genomes from both colonies, we used VIBRANT[56] for prophage prediction and dRep [57] to group phage sequences into related clusters. With this method, we identified 91 prophages of sufficient quality, which were then clustered into seven genus-level clusters (80% ANI), five of which are found in *B. apis* genomes isolated from our colonies (Fig. 4C). Sixty percent of phage clusters are found in both colonies (Fig. S4A, C), with two clusters (2 and 4) unique to colonies D and J respectively. To examine phage distribution at a finer genetic resolution, we grouped phages with at least 95% ANI into subclusters (34 distinct subclusters). Very few subclusters (23.5%) were identified in both colonies (Fig. S4B). Rather, most phage subclusters were unique to a single colony (76.5%), and in many cases, unique to one *B. apis* genome (Fig. S4D). Together, this suggests that although phage clusters are often shared between colonies, rapid diversification of phages may lead to distinct subclusters, unique to colonies.

**Figure 4:**
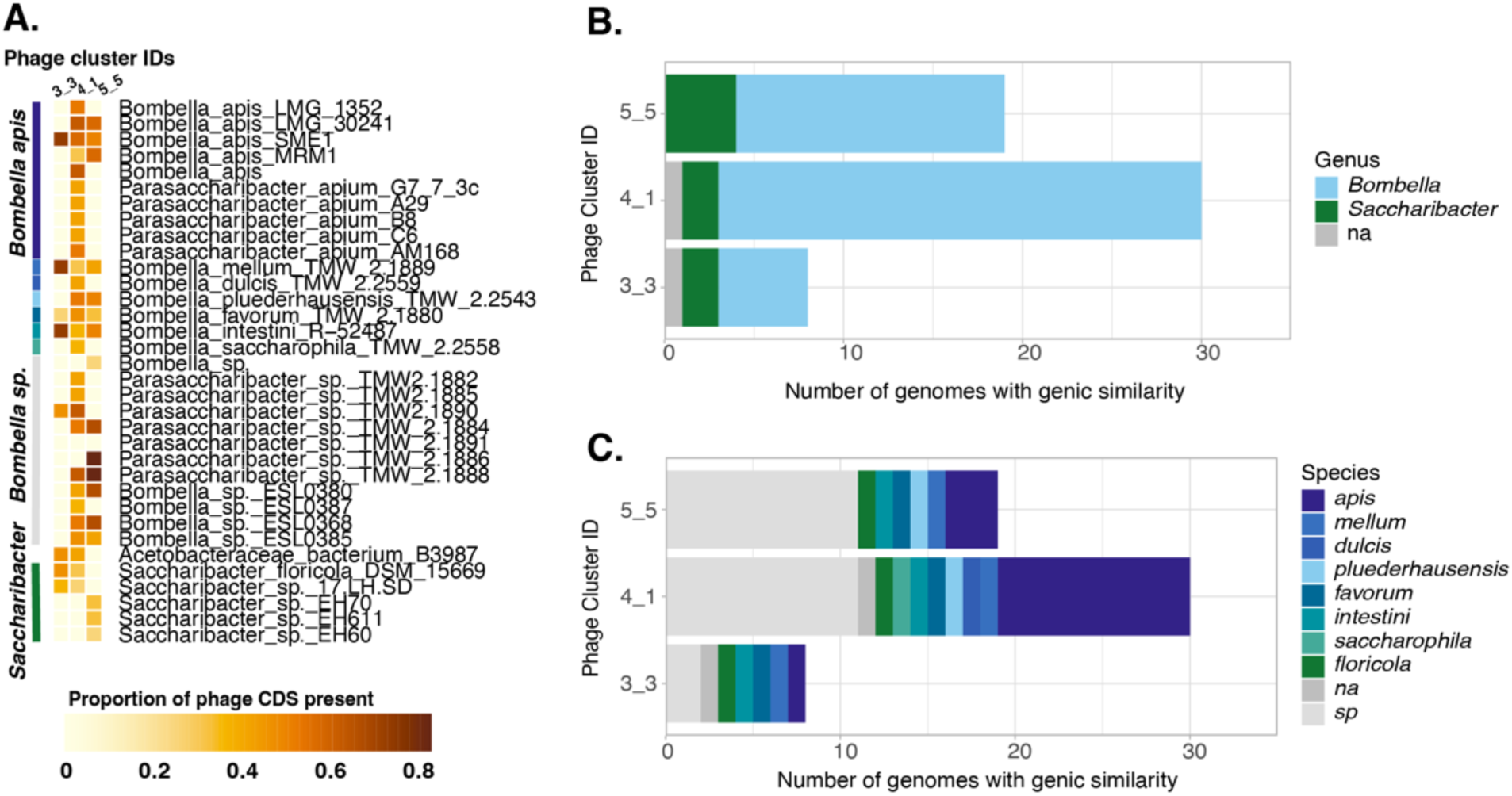
*B. apis* phages have high genic similarity with CDSs from other *Bombella* species and *Saccharibacter* genomes. (A) The proportion of phage-encoded genes present in bacterial genomes. One representative phage from each cluster was used as a tBLASTn query against all publicly available genomes on NCBI, requiring 50% query coverage and percent amino acid identity. Only three representative phages returned hits. To determine if CDSs were syntenic -as would be expected for phage-encoded genes -cblaster was used, requiring at least 25% of phage encoded genes to be present in a neighborhood less than or equal to the phage length* 1.5 to allow for phage size expansion. If all the above criteria were met, the bacterial genome was considered a hit. The proportion of query phage proteins present in each hit genome is shown in the first panel (A). For each phage cluster that returned hits, the hits were classified by the (B) genus and (C) species of its host genome assembly. Assemblies with no designation at the genus level are labeled ‘na’ and those without designated species are labeled ‘sp’.

### Similarity of *B. apis* phages to CDSs from other *Bombella* and *Saccharibacter* strains

To determine if *B. apis* may be acquiring phages from other bacterial species, either other honey bee symbionts or environmental bacteria, we searched for any bacterial genomes with high amino acid similarity to phages identified in our colonies. Querying all publicly available genomes on NCBI using cblaster [51], we identified gene clusters with at least 50% amino acid identity and 50% query coverage to representative phages from each phage cluster. Three of seven phage clusters returned hits (Fig. 4A), suggesting that either phages or phage-encoded cargo genes are shared with these genomes. All hits were to AAB, specifically *Bombella* and *Saccharibacter*, a clade of flower- and solitary bee-associated AAB sister to *Bombella* [61–63] (Fig 4B), and most hits were from *B. apis* or other *Bombella* species isolated from honey or bumble bees (Fig. 4C). What functions the shared CDSs encode is unclear, given that most sequences shared between phages and cblaster hits are of unknown function (Fig. S3).

### Identification of plasmids from nectar, nurse crop, and queen gut isolates

During genome assembly, we identified five closed extrachromosomal elements in four genome assemblies (AML2, JPL21, JPL24, and SME1) (Fig. 5B). To determine if these closed contigs were plasmids we manually annotated with BlastX to identify replication machinery. In all five extrachromosomal elements, replication proteins were present (Fig. 5A), belonging to either RepA, RepB, or RepC families (Table S1). Additionally, we visualized DNA fragments of approximately the correct length in plasmid preps from each strain (Figure S2). Interestingly, all plasmids were found in isolates from nectar, nurse crops, and queens (Fig. 5A), but never in larval isolates (Figure 5A). These are the first sequenced plasmids from *Bombella*, although it is likely that short-read sequencing may be obscuring plasmids in older assemblies. To quantify overall nucleotide similarity between plasmids, we measured Jaccard similarity with sourmash [53] . Two pairs of plasmids (pJPL24_01 and pSME1; pAML2 and pJPL21) are similar, with a Jaccard Index of 0.1 (Fig. 5B). Further annotation of these plasmids revealed that three of five plasmids have conjugation machinery, suggesting a possible HGT mechanism.

**Figure 5:**
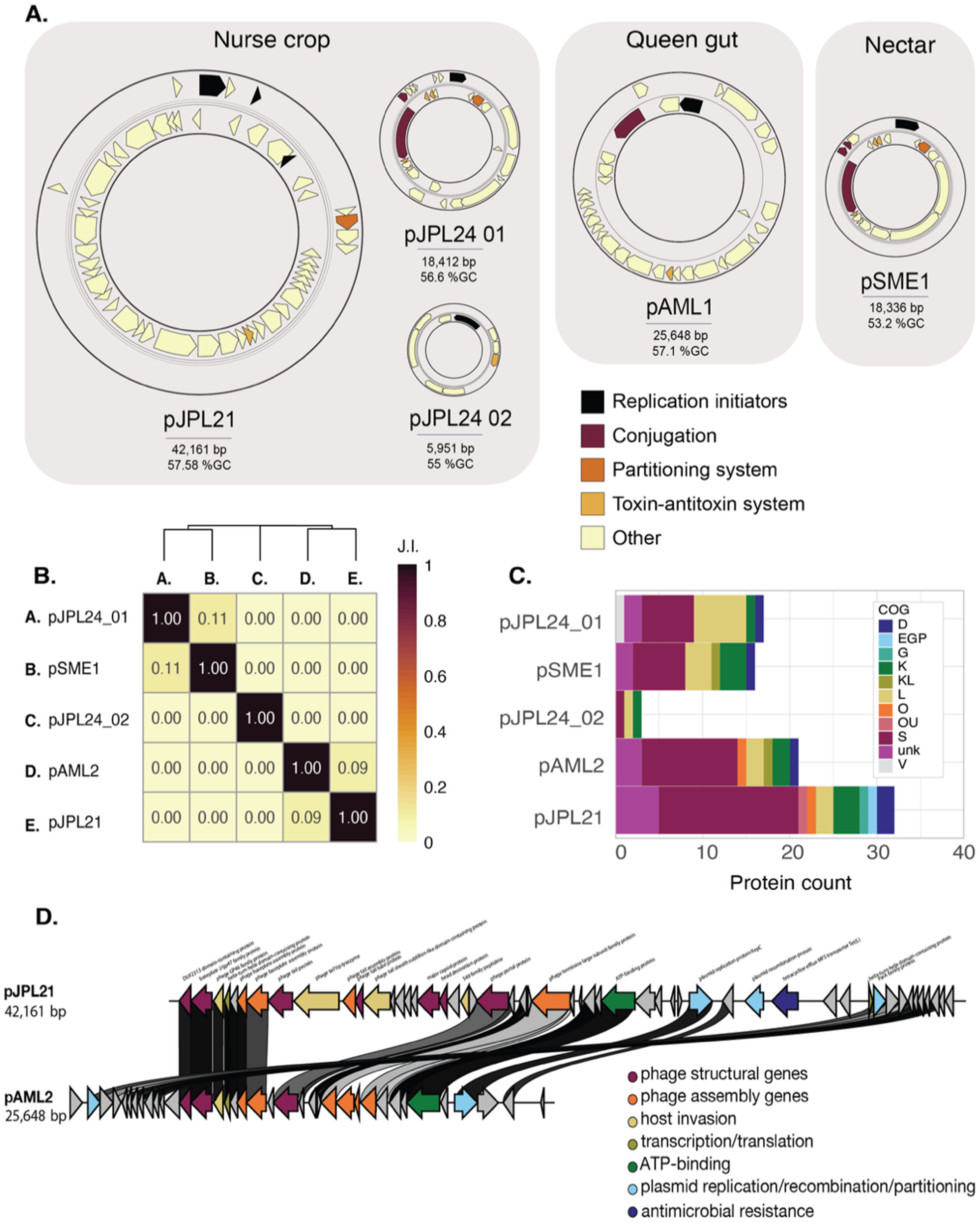
Five unique plasmids were identified in isolates from nectar, nurse crops, and queen guts. (A). Four strains harbored plasmids (JPL21, JPL24, AML1, SME1). Plasmid replication machinery and cargo genes are colored by annotation. (B). Overall nucleotide similarity of *B. apis* plasmids to one another was measured with Jaccard similarity using sourmash. Scores above 0.1 indicate that plasmids are significantly similar. (C) COG categories of plasmid-encoded genes. Plasmid coding sequences were annotated by eggNOG. Colors denote COG functional categories. (D) Alignment of putative phage-plasmids pJPL21 and pAML2 showing percent amino acid similarity of phage and plasmid genes as annotated by Pfam.

### Plasmids encode toxin-antitoxin systems, antimicrobial resistance cassettes, and prophage regions

To gain insight into the function of the plasmid cargo genes, we annotated them with eggNOG (Table S2) in addition to PGAP. Most proteins were not classified by COG categories (Fig.4C). Of these unclassified proteins in pAML2 and pJPL21, many had high homology to phage proteins in Pfam and annotation with VIBRANT confirmed the presence of similar phages genes on both pAML2 and pJPL21. Since both encode plasmid replication proteins in addition to this phage, pAML2 and pJPL21 would more appropriately be classified as phage-plasmids (P-Ps). When compared to phages identified in *B. apis* genomes, both P-Ps were placed in cluster 1, a cluster not otherwise represented in our colonies (Fig. 3C). In addition to sharing similar phages, both P-Ps have high genetic similarity (Jaccard index: 0.09) (Fig. S6B); the larger of the two, pJPL21, encodes 86% of genes found on the smaller P-P, pAML2, but has a stretch of approximately 17,000 bps unique to it, including a tetracycline resistance cassette (Fig.S6A). In short, both P-Ps are similar to each other, but otherwise bear no similarity to other phages or plasmids found within our colonies, instead clustering with phages found in *Bombella* isolates from Europe. One unifying functional feature across all plasmids was the presence of toxin-antitoxin (T-A) systems (Figure 5A; Table S1). All T-A systems identified are Type II, producing a stable toxin protein and an unstable antitoxin. Likely these systems are involved in plasmid maintenance but could also play a role in phage defense [64].

### Plasmid coding sequences share similarity with acetic acid bacteria (AAB), including other *Bombella apis* strains

Since plasmids are mosaic, and frequently take up and lose their cargo genes, taxonomically classifying plasmids based on sequence similarity is challenging. To approach this classification issue, we took two different approaches: (1) one to find syntenic regions with similar coding sequences to our plasmids and (2) to identify plasmids with similar backbones. The first approach repurposed software developed for biosynthetic gene clusters to identify contiguous clusters of coding sequences (CDS) with high similarity to CDSs from our plasmids. This approach has the advantage of identifying clusters of similar CDSs regardless of whether they are located on chromosomes, plasmids, etc. The second approach used nucleotide similarity across short regions (k=31 bp) to identify closely related sequences. This approach is more stringent and has a higher chance of recovering plasmid sequences with the same backbone, or non-coding sequences.

Using the first approach, we identified similar gene clusters in AAB and in one LAB from honey bees (Figure 6A). *Acetobacter*, *Gluconobacter*, and *Komagaetibacter* CDSs cluster with three of five plasmids. On average, around half the CDSs encoded on each plasmid were found in other genomes. The *B. apis* plasmids pJPL24_01, pJPL24_02, and pSME1 all had similar CDSs from known plasmids (Figure 6A and B). On the other hand the P-Ps, pAML2 and pJPL21, only shared similar CDSs with other *Bombella* (formerly *Parasaccharibacter*) and *Saccharibacter* genomes (Figure 6A and B). Two *Bombella* genome assemblies (LMG1352 and TMW 2.1884) shared 91% and 88% of pAML2’s CDSs respectively. *Saccharibacter* assemblies (EH611, EH60 and EH70) by comparison shared only 55-58% of pAML2’s CDSs (Figure 6A). All hits to our plasmids were either AAB or LAB and isolated from acidic environments such as insect guts, nectar, or fermented foods (Figure 6C). P-Ps, pAML2 and pJPL2, were most similar to other bacterial genomes isolated from honey bees, whereas all other plasmids were most similar to AAB isolated from fermented food, fruits, flowers, and plants (Figure 6C).

**Figure 6:**
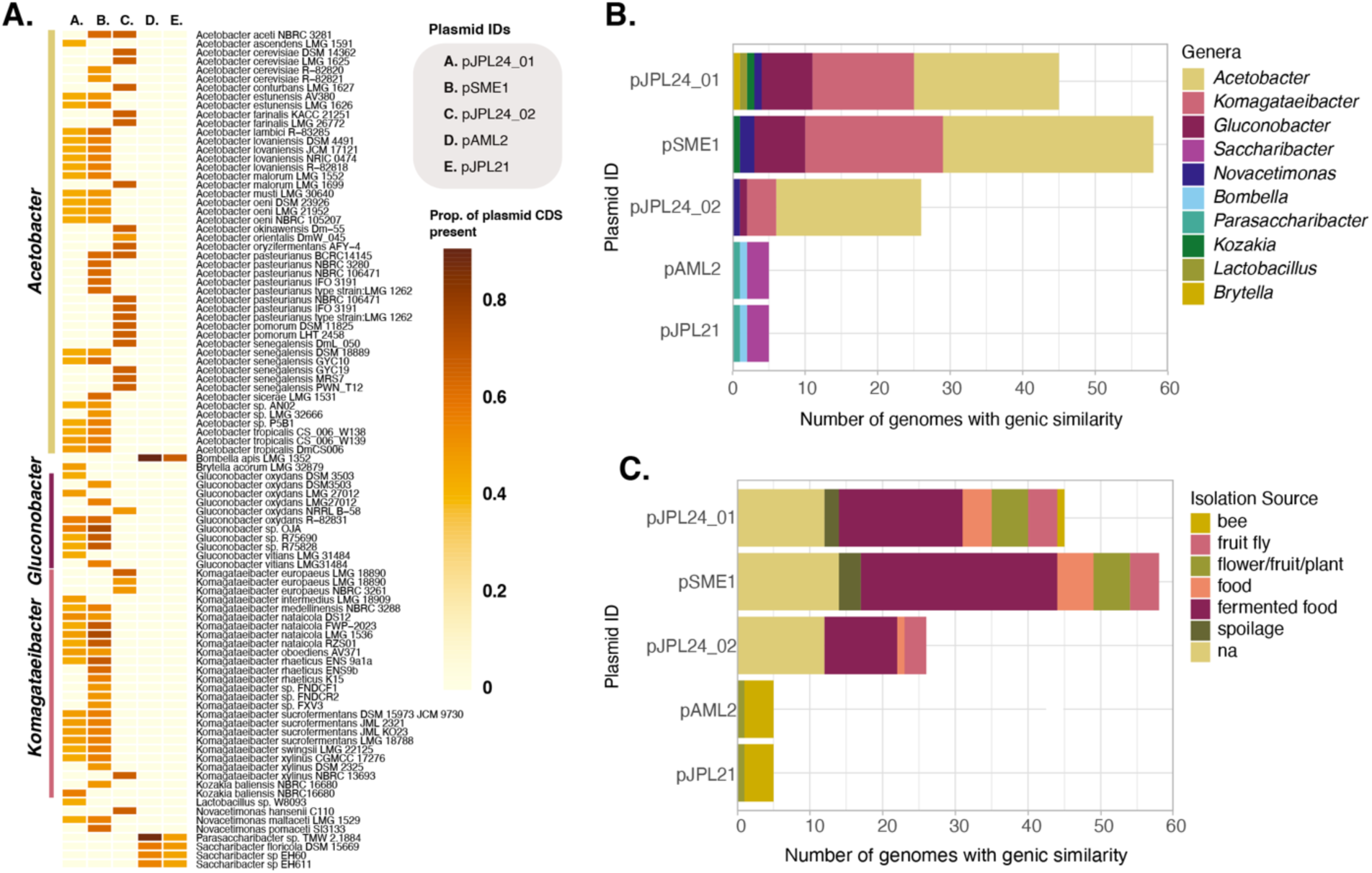
Plasmid-encoded genes have high similarity to CDS found among other acetic acid bacteria isolated from insects, flowers, and fermented foods. (A) The proportion of plasmid-encoded genes present in bacterial genome assemblies. The presence of CDSs with high similarity to *B. apis* plasmid CDSs in all NCBI genomes was determined using tBLASTn with 50% query coverage and 50% amino acid identity. To determine if CDSs were present in clusters instead of dispersed across the genome, cblaster was used, requiring that at least 25% of plasmid CDSs were present within a range equal to the (plasmid length) * 1.5 to allow for plasmid expansion. If all the above criteria were met, the assembly was considered a hit. (B) Plasmid CDS hits were characterized by genera or (C) isolation source as reported on NCBI.

### Plasmid replication and conjugation machinery is highly conserved among a small number of AAB plasmids

Our second approach to identify similar plasmids quantified overall nucleotide sequence similarity, which encompasses both the plasmid backbone and coding regions. Using sourmash, plasmids were subset into k-mers of 31 bps, and pairwise nucleotide similarity, as defined by Jaccard Similarity, was calculated. We defined similar plasmid backbones as having a Jaccard Index ≥ 0.1. This threshold has previously been used to classify similar plasmids into “clusters” or “cliques” [65, 66]. Using these criteria we queried three different databases of plasmid sequences. First, we queried all the known plasmids that have been isolated from honey bee symbionts and pathogens and are available in the Plasmid Database (PLSDB) but found no hits (Figure S5). Secondly, we created a database of plasmid sequences identified as hits during our CDS cluster similarity analysis (Figure 6A). Five plasmids from this database had a Jaccard index value greater or equal to 0.1, indicating similarity (Figure 7A). Thirdly, we built a database of plasmid sequences with similarity to our plasmid’s replication proteins (top 100 hits from a tBlastN query). Sequence similarity analysis of this database yielded three new hits with significant Jaccard index values and one hit already identified by CDS cluster similarity (*Komagataeibacter nataicola* strain DS12 plasmid pKNA10) (Figure 7A). Replication machinery was highly conserved between pJPL24_02 and its top hit, pKHC110_1 (Figure 7B). pSME1 followed a similar trend, sharing strong homology with the RepA protein and recombinase from its top hit, 2P (Figure 7C). In the case of pJPL24_01, however, a large region (∼6 kb) is highly conserved with pKNA10, a plasmid from *Komagataeibacter nataicola*. This region encodes RepA, conjugal transfer proteins, TraD and TraA, as well as a VapB antitoxin (Figure 7D). Together this suggests that three *B. apis* plasmids isolated from our apiary possess backbones, replication machinery, and some cargo genes homologous to known plasmids in AAB from fermented foods, but are not similar to any known plasmids from honey bee-associated bacteria.

**Figure 7:**
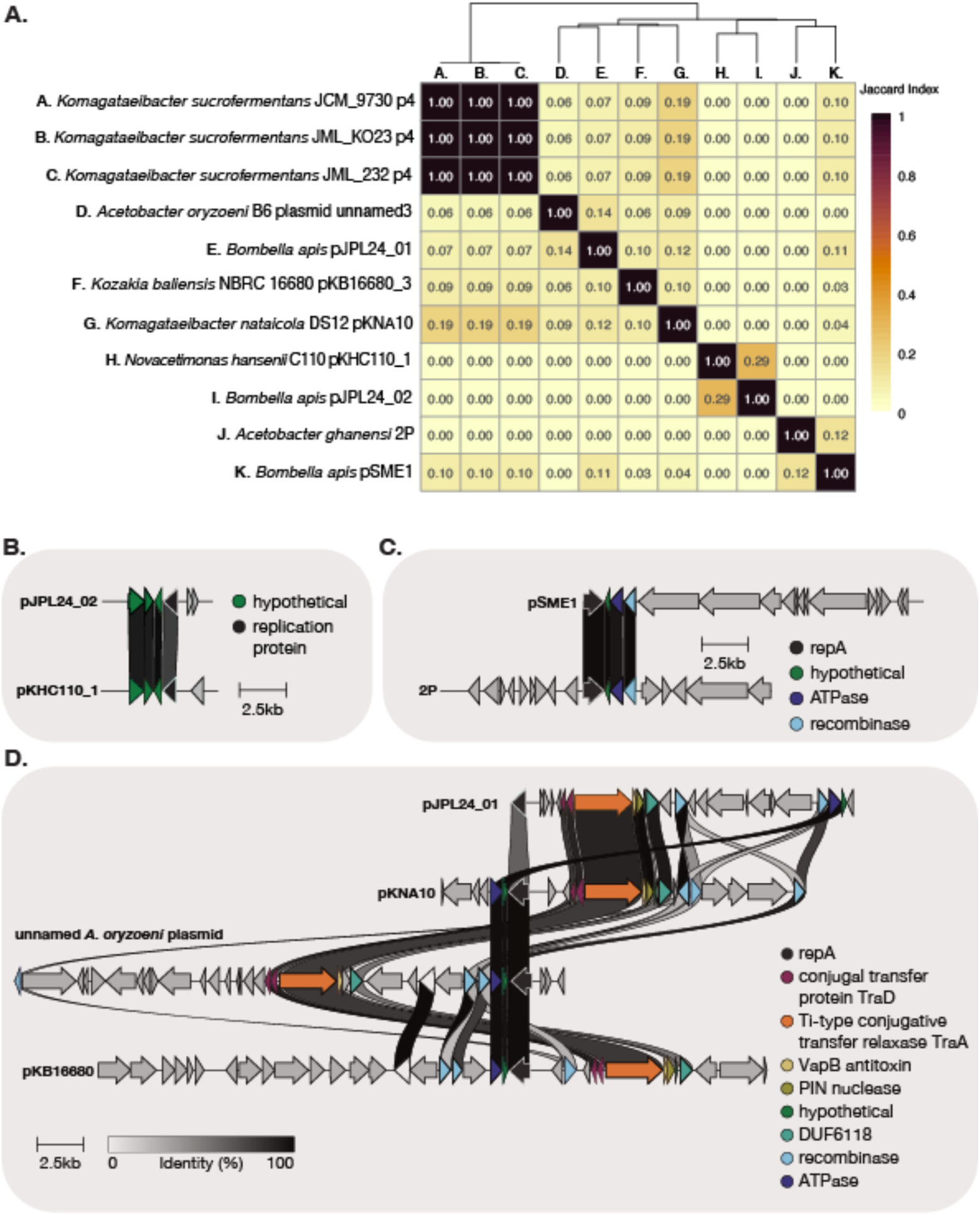
*B. apis* plasmids have low overall similarity to plasmids from other AAB but have highly conserved replication and conjugation regions compared to AAB plasmids. (A) Similarity of plasmids to other AAB plasmids as determined by Jaccard-Index. Alignment of conserved plasmid regions from (B) pJPL24_02, (C) pSME1, and (D) pJPL24_01. Colors of genes represent PGAP functional annotation. Unaligned genes are shown in gray. Pairwise amino acid identity between CDS is shown in greyscale.

## Discussion

The Acetobacteraceae *B. apis* is an important symbiont of honey bees – both buffering against poor nutrition and reducing the likelihood of fungal infections during larval development [17, 18]. The diversity of *B. apis* strains within and between colonies and their distribution across colony environments has not previously been examined, since most of our understanding o*f B. apis*’ ecology is based on 16S rRNA amplicon studies that lack the resolution to study strain variation [33, 34, 38]. Here we isolated and sequenced nineteen new *B. apis* strains and identified MGEs, plasmids, phages, and phage-plasmids (P-Ps) that contribute to their genetic diversification. Additionally, we used two complementary approaches to identify MGE relatives in other bacteria, with the goal of identifying possible sources from which *B. apis* acquired MGEs.

In this process we identified and isolated two strains of *Bombella favorum*, a close-relative of the bumblebee symbiont, *Bombella intestini*. This finding, in corroboration with prior studies, suggests that other *Bombella* species are present in the colony [60]. Whether this association is transient or perpetual is unclear. Given the high similarity of the 16S rRNA operon between *Bombella* species, it is possible that OTUs ascribed to *B. apis* in past studies actually represent a consortium of *Bombella* species, but further studies are needed to determine their association with honey bees.

Within the *B. apis* clade there is no striking correlation between geography and genetic relatedness. Genetic variation in other symbiont populations, such as those associated with bobtail squid and deep sea snails, is largely explained by geographic location [67, 68]. This striking lack of correlation is perhaps representative of honey bee management practices. The European honey bee is, after all, a domesticated insect, having been managed by humans for 4,000 years and transported across continents for its pollination services [69, 70]. New colonies, which assemble naturally by swarming, are now generally founded by beekeepers with queens bred in queen-breeding facilities [71]. In the US, only a handful of queen-breeding companies supply most commercial and private beekeepers. Given *B. apis’* association with the queen gut, it seems likely that queen-breeding facilities serve as reservoirs, and that colonies across the US are seeded with strains derived from these queens.

Strains derived from our colonies did not cluster by their isolation environment: larvae, queens, nurse crops, or nectar. Instead, larval isolates are clustered with strains from all other hive environments. The bias in our sampling towards larval strains impedes our ability to statistically test for correlation between isolation environment and phylogenetic relatedness and draw a more meaningful conclusion. To test for spatial segregation or niche partitioning of strains (as observed in other symbionts[72–75] within the hive, both deeper sampling and longitudinal data are needed. The clearest distinction between larval and non-larval strains was the distribution of plasmids among them. Despite our larger sampling of larval strains, none carried any detectable plasmids, but all other strains had at least one plasmid. Based on this observation, and the fact that geography can dictate plasmid transmission and evolution [76], it may be worthwhile to examine whether *B apis*’ spatial distribution within the colony correlates with plasmid abundance and distribution in future studies.

Plasmid transmission and evolutionary history in natural microbial communities are still poorly understood. Due to the mosaic composition of plasmids, and other mobile genetic elements, we need different evolutionary frameworks to disentangle their patterns of diversification [24, 76]. To search publicly available genomes for similar plasmids or plasmid cargo genes, we adopted a tool, cblaster, developed for other horizontally transferred genes: biosynthetic gene clusters [51]. This method allowed us to survey draft genome assemblies in an unbiased fashion. Some of our hits were also confirmed by our alternative method (Jaccard similarity) when used on a subset of known plasmid sequences. Whether the CDS present in other AAB assemblies are chromosomal or on plasmids is not known but would be an interesting avenue of future study. The coding sequences present in other AAB are largely unannotated (Figure 4C), but their prevalence across diverse AAB may be indicative of a function important for this phylogenetic group. Alternatively, if these CDSs are localized to plasmids, their distribution throughout AAB may be the result of the plasmid’s broad host range and horizontal transfer.

How did *Bombella* acquire these plasmids? The strong similarity in both synteny and gene content between the two P-Ps and other *B. apis* and *S. floricola* assemblies suggests that highly similar P-Ps may be stably associated with honey bees. The other three plasmids, however, have low similarity to plasmids from AAB isolated from plants and fermented foods. Typically, CDSs shared between *B. apis* and AAB plasmids were involved in replication, recombination and conjugation and did not include cargo genes carried on the plasmid backbone. Perhaps the similarities between them are indicative of a common plasmid backbone with a broad host range, that has infected AAB sporadically throughout their evolutionary history.

Likely, *B. apis* acquired these three plasmids from a plant-associated AAB, but whether vertically or horizontally is challenging to disambiguate. Both insect- and fermented food associated-AAB diverged from a plant-associated clade [36]. Therefore the similarity of *Bombella*’s plasmids to genera associated with flowers, namely *Neokomagataea* [77, 78], *Gluconobacter*, and *Saccharibacter* [62] could be indicative of an ancient association or a recent acquisition. In the latter scenario, flowers may act as transmission hubs for the exchange of AAB symbionts and their MGEs [35]. This phenomenon has been documented for pollinator pathogens, which can swap hosts via shared floral resources [79]. Regardless of how *B. apis* acquired these plasmids, it is clear that they belong to a group of similar plasmids that can be harbored by AAB across diverse environments.

Most studies of MGEs in honey bees focus on the gut microbiome of adult workers -which is compositionally distinct from the communities in which *B. apis* resides. Connecting our findings to these studies of phages and plasmids in the adult microbiome, however, reveals some commonalities in the distribution of MGEs within and across colonies. In the adult gut, plasmids and phage clusters are shared between different strains and species within the same colony, as well as across colonies in the same apiary [24]. For phages, the same clusters can be identified in bacterial genomes from colonies worldwide, suggesting a widespread association. It is likely that some of these phages are also P-Ps, given that many also encode RepC proteins. In the adult worker microbiome, phages often encode metabolic genes with obvious links to host nutrition, but the same is not true for *B. apis* phages [25]. Future studies interrogating the function of the phage cargo genes could elucidate their role in shaping *B. apis*’ phenotypic diversity.

Overall, this study has identified novel phages, plasmids and phage-plasmids associated with the important honey bee symbiont *B. apis* and documented their distribution across colonies and colony environments. Our work highlights the role of phages in driving genetic diversity between *B. apis* strains within the same colony and apiary - a finding reflected in studies of other honey-bee microbiome members. In contrast, the plasmids and P-Ps we identified were rare within our sampling and only found in a few strains isolated from queens, nurses, and food stores. While both P-Ps clustered with other *B. apis* phages, the plasmids had low genic similarity to AAB plasmids from plants and fermented foods. To better interpret these findings, *B. apis*’ transmission within and between colonies should be explored, as well as the influx of AAB into the colony. The interconnectedness of AAB populations, linked by the trophic interactions of their hosts, represents a fascinating area of study for tracing the transmission of mobilized traits across different symbiont populations.

## Notes

### Competing Interest Statement

The authors have declared no competing interest.

### Funding

- NSF: NSF DBI 2022049
- USDA: 2022-67011-36577

