## Supplemental Figures and Tables for "Nested Connections: Local Phage and Broad Plasmid Sharing in the Honey Bee Mobilome"

### Supplementary Figures

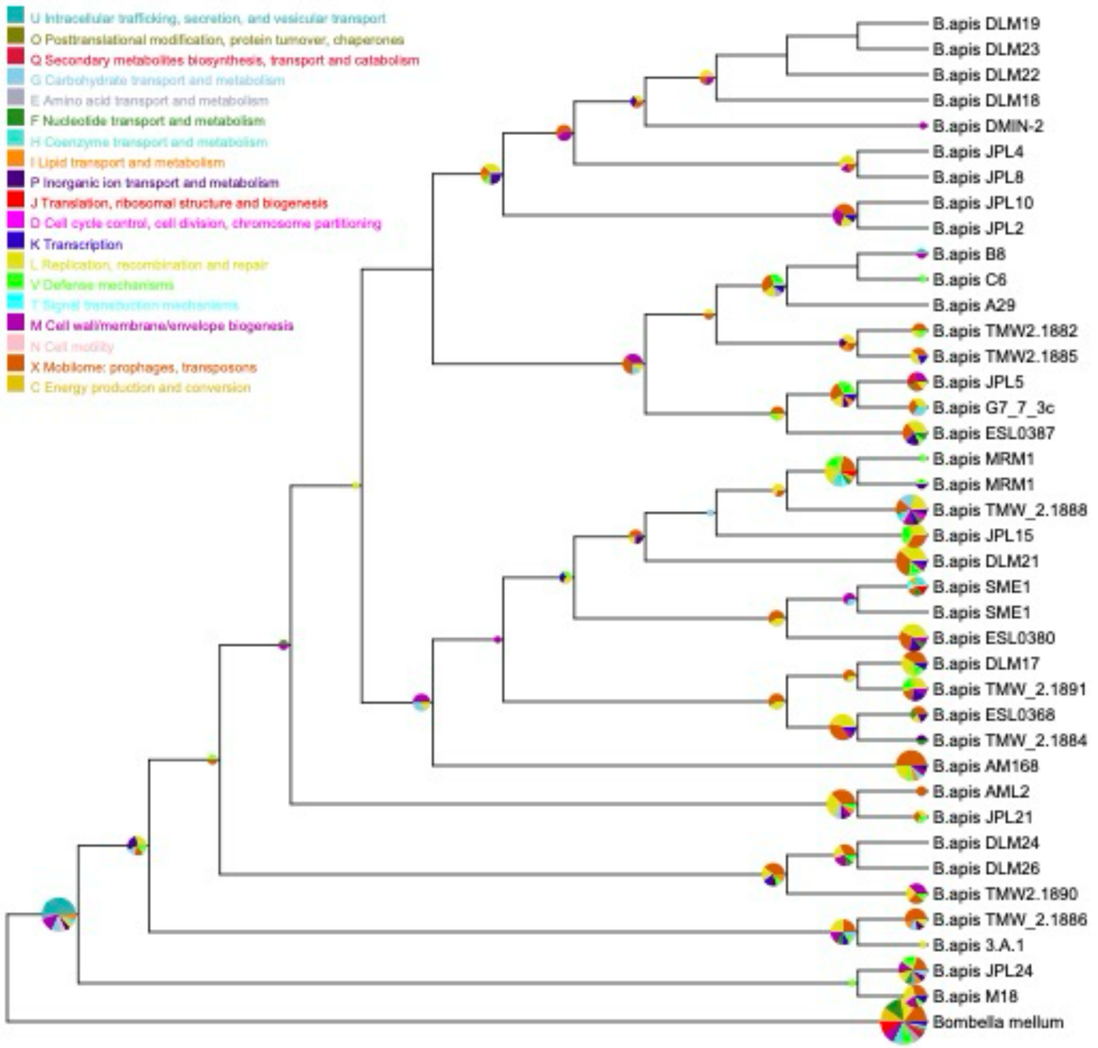


**Figure S1: Gene losses with the *B. apis* clade.** COG categories of genes lost within the *B. apis* clade. Gene gains and losses were defined with GLOOME and categorized by their COG family. Pie charts denote the COG categories of genes lost at that node or leaf. Pie chart colors represent COG categories (U: intracellular trafficking, secretion and vesicular transport; O: posttranslational modification, protein turnover and chaperones; Q: secondary metabolite biosynthesis, transport, and catabolism; G: carbohydrate transport and metabolism; E: amino acid transport and metabolism; F: nucleotide transport and metabolism; H: coenzyme transport and metabolism; I: lipid transport and metabolism; P: inorganic ion transport and metabolism; J:translation, ribosomal structure and biogenesis; D: cell cycle control, cell division, and chromosome partitioning; K: transcription; L: replication, recombination, and repair; V: defense mechanisms; T: signal transduction mechanisms; M: cell wall/membrane/envelope biogenesis; N: cell motility; W: extracellular structures; X: mobilome, prophage, and transposons; C: energy production and conversion) and the diameter of each pie represents the number of genes gained.

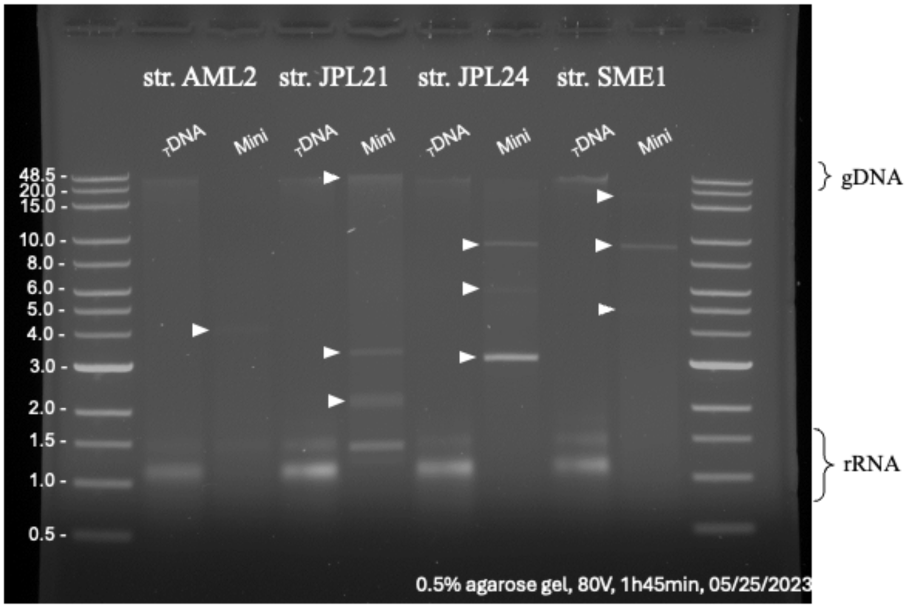


**Figure S2: Plasmid preps and gDNA from *B. apis* strains.** Total DNA and plasmid extractions from B. apis strains with putative plasmids, visualized by gel electrophoresis. Approximately 150 ng of DNA was visualized on a 0.5% agarose gel run for 100 minutes at 80 Volts.

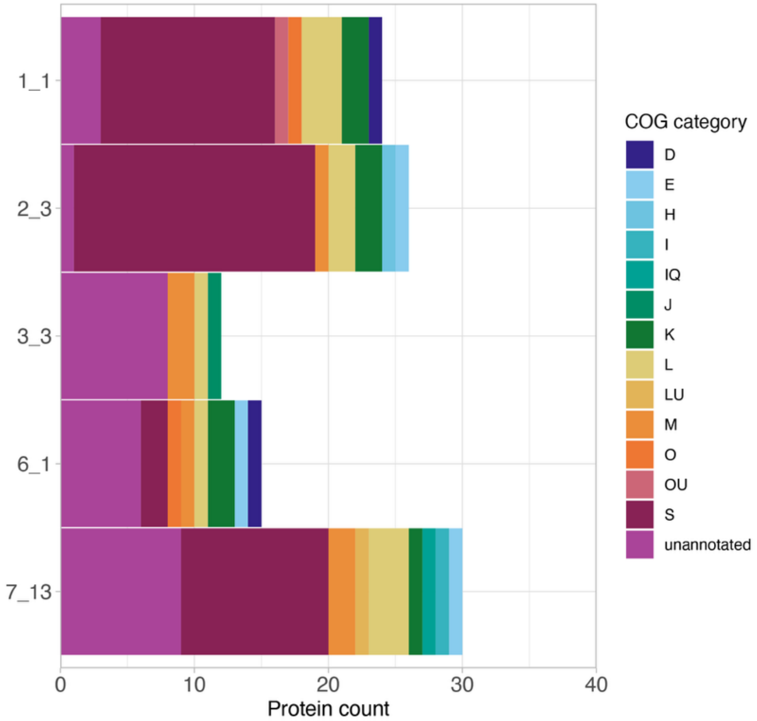


**Figure S3: Functional categorization of phage-encoded genes.** COG categorization of *B. apis* phage-encoded genes. Phages identified by VIBRANT were annotated by eggNOG to classify each CDS by its COG category. The majority of phage cargo genes – genes not involved in the replication and maintenance of the phage – are unclassified.

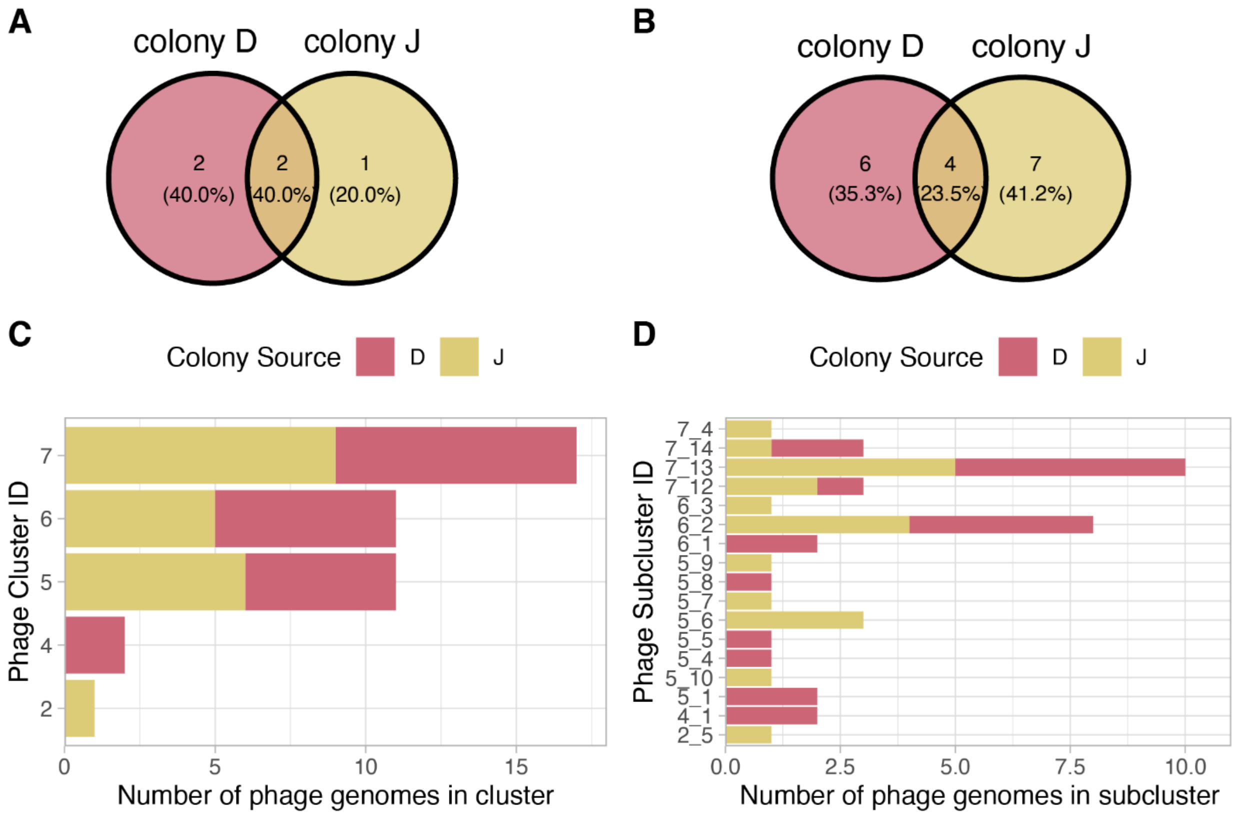


**Figure S4: Phage subclusters are frequently found in only one colony OR Phage clusters are frequently shared between colonies.** The presence/absence of phages in *B. apis* strains isolated from either colony D or J at our apiary. Phage clusters are composed of phages sequences with 80% ANI with 80% query coverage, and subclusters are composed of phages with 95% ANI with 80% query coverage. (A) Representatives from most phage clusters are found in both colony D and J. (B) The absolute abundance of phages grouped by their cluster and colony source. Clusters found in both colonies were also the most abundant. Clusters unique to only one colony were relatively rarer within that colony. (C) At a finer resolution, phage subclusters are largely unique to one colony or the other. (D) The absolute abundance of phages grouped by their subcluster and colony source. Subclusters found in both colonies were more abundant, whereas subclusters found in only one colony were often unique to only one *Bombella* strain.

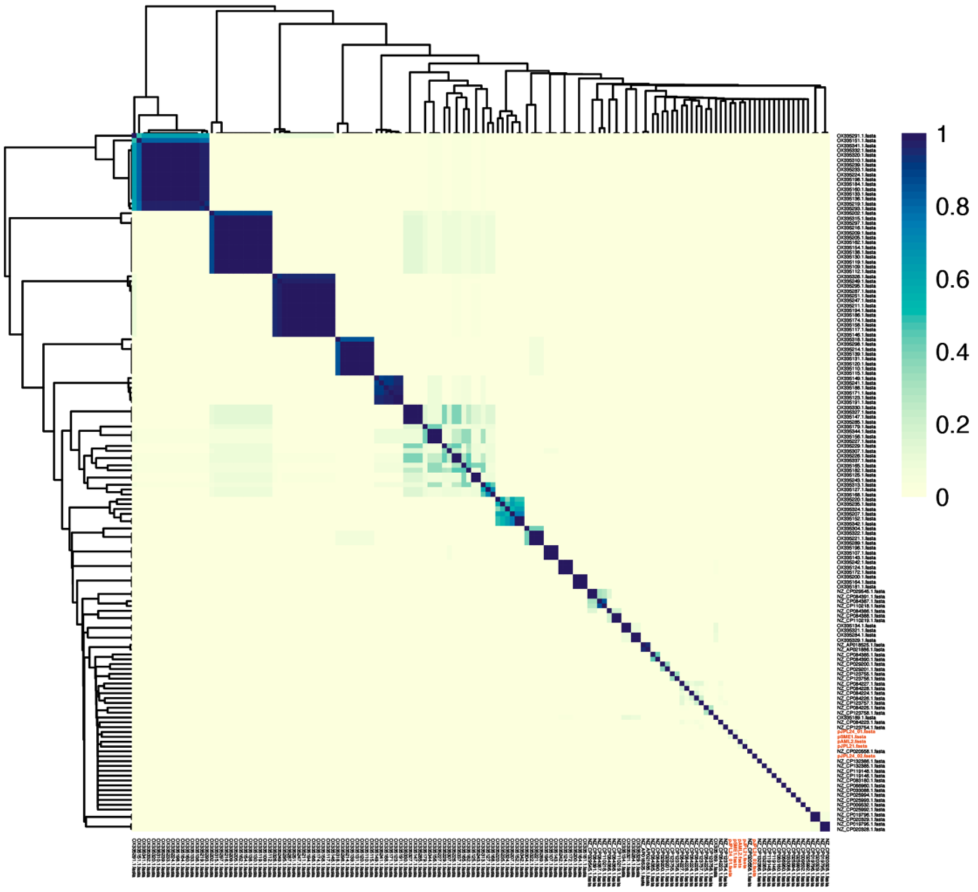


**Figure S5: *B. apis* plasmids bear low similarity to other plasmids isolated from honey bee colonies** Overall nucleotide similarity of *B. apis* plasmids to plasmid database (PLDB) plasmids isolated from honey bees was measured with Jaccard similarity using sourmash. Scores above 0.1 indicate that plasmids are significantly similar. Plasmids from this study are highlighted in red.

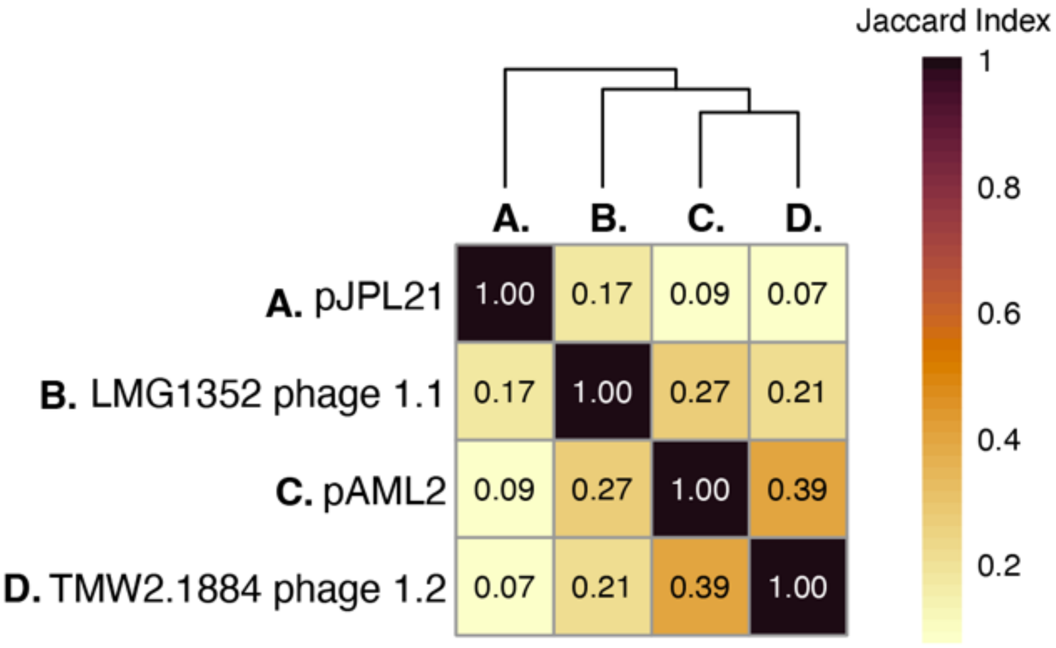


**Figure S6: Putative phage-plasmids (P-Ps) isolated from queens and nurse crops are significantly similar to phages identified in other *B. apis* strains** Overall nucleotide similarity of *B. apis* P-Ps to phages identified in other *B. apis* strains belonging to cluster 1 was measured with Jaccard similarity using sourmash. Scores above 0.1 indicate significant similarity.

### Supplementary tables

**Table S1: Plasmid manual annotation.** Genes of interest in plasmid assemblies were annotated manually with BLASTx. Replication proteins, conjugation machinery, partitioning systems, and toxin/anti-toxin systems were focused on due to their putative role in replication, transmission, and persistence of the plasmids.

| | **pAML2** | **pJPL21** | **pJPL24_01** | **pJPL24_2** | **pSME1** |
|---|---|---|---|---|---|
| Replication initiator | *repC* | *repA* | *repA* | *repB* | *repA* |
| | (AML2p1_000001; MCL1512564) | (JPL21p1_000004; EHT8928499) | (JPL24p1_000001; WP_048848528) | (JPL24p2_000001; GBR53535) | (SME1p1_000001; KXV21260) |
| | | *repB* | | | |
| Conjugation | | (JPL21p1_000007; EGP5572984) | | | |

| | | | | | |
|---|---|---|---|---|---|
| | | *repC* | *trA* | | Ψ*traA* |
| | | (JPL21p1_000001; WP_282803146) | (JPL24p1_000018; WP_173575238) | | (SME1p1_000012; WP_012813389) |
| Partition | | | *traD* | | *traD* |
| Toxin/Anti-toxin | *parA* | | (JPL24p1_000018; WP_173575238) | | (SME1p1_000013; WP_173575238) |
| | (AML2p1_000030; MCL1512568) | | *traD* | *relE*/*parE* | *traD* |
| | *relE* | *parA* | (JPL24p1_000019; WP_086442351) | (JPL24p2_000004; MBS0990063) | (SME1p1_000014; MCT6837494) |
| | (AML2p1_000015; WP_249080617) | (JPL21p1_000011; WP_249080663) | *parA*/*minD* | | *parA* |
| | | *relE*/*parE* | (JPL24p1_000003; WP_010511349) | | (SME1p1_000003; WP_206035346) |
| | | (JPL21p1_000025; WP_249080617) | *vapB* | | *vapB* |
| | | | (JPL24p1_000017; WP_010516209) | | (SME1p1_000011; WP_199283227) |
| | | | *brnA* | | *brnA* |
| | | | (JPL24p1_000023; WP_086442348) | | (SME1p1_000017; WP_034958702) |
| | | | *brnT* | | *brnT* |
| | | | (JPL24p1_000024; WP_063907132) | | (SME1p1_000018; MCT6837492) |
| (Ψ) pseudogene | | | | | |

**Table S2: Plasmid eggNOG annotation.** All CDSs in plasmid assemblies were annotated with eggNOG-mapper to classify them by COG functional category and Pfam domains.

| plasmid | evalue | score | COG_category | Description | PFAMs |
|---|---|---|---|---|---|
| pSME1 | 1.75e-254 | 699.0 | K | Filamentation induced by cAMP protein fic | Fic,Fic_N |
| pSME1 | 1.68e-149 | 421.0 | - | - | - |
| pSME1 | 5.03e-12 | 60.5 | S | Plasmid stability protein | Arc |
| pSME1 | 3.01e-84 | 249.0 | S | Toxic component of a toxin-antitoxin (TA) module. An RNase | PIN |
| pSME1 | 5.5e-34 | 117.0 | K | Bacterial antitoxin of type II TA system, VapB | VapB_antitoxin |
| pSME1 | 5.18e-64 | 197.0 | S | Conjugal transfer protein TraD | TraD |
| pSME1 | 6.14e-53 | 166.0 | S | Conjugal transfer protein traD | TraD |
| pSME1 | 5.71e-53 | 166.0 | S | CopG antitoxin of type II toxin-antitoxin system | CopG_antitoxin |
| pSME1 | 2.93e-56 | 175.0 | S | Ribonuclease toxin, BrnT, of type II toxin-antitoxin system | BrnT_toxin |
| pSME1 | 2.52e-243 | 671.0 | L | Replication initiator protein | RPA |
| pSME1 | 5.96e-43 | 142.0 | - | - | - |
| pSME1 | 4.44e-142 | 402.0 | D | VirC1 protein | AAA_31,CbiA,VirC1 |
| pSME1 | 0.0 | 1177.0 | KL | Type III restriction enzyme, res subunit | ResIII |
| pSME1 | 0.0 | 928.0 | L | PFAM DNA methylase N- | N6_N4_Mtase |

| | | | | | |
|---|---|---|---|---|---|
| | | | | 4 N-6 domain protein | |
| pSME1 | 1.4e-131 | 374.0 | L | Resolvase, N terminal domain | Resolvase |
| pSME1 | 1.06e-95 | 280.0 | K | Transcriptional regulator | ROS_MUCR |
| pAML2 | 6.94e-212 | 602.0 | S | COG5511 Bacteriophage capsid protein | Phage_portal_2 |
| pAML2 | 5.01e-37 | 136.0 | S | DNA circularisation protein N-terminus | DNA_circ_N |
| pAML2 | 1.72e-150 | 435.0 | S | Mu-like prophage tail protein | Phage_GPD |
| pAML2 | 5.85e-81 | 244.0 | S | Bacteriophage Mu Gp45 protein | Phage_Mu_Gp45 |
| pAML2 | 1.04e-22 | 90.9 | K | Helix-turn-helix domain | HTH_3 |
| pAML2 | 3.97e-57 | 183.0 | S | Phage protein GP46 | GP46 |
| pAML2 | 1.77e-181 | 514.0 | S | Baseplate J-like protein | Baseplate_J |
| pAML2 | 4.9e-67 | 211.0 | S | Tail protein | DUF2313 |
| pAML2 | 1.05e-14 | 80.5 | - | - | - |
| pAML2 | 8.98e-52 | 167.0 | - | - | - |
| pAML2 | 5.35e-117 | 338.0 | D | plasmid maintenance | AAA_31 |
| pAML2 | 7.03e-276 | 793.0 | KL | HELICc2 | Helicase_C_2,ResIII |
| pAML2 | 1.49e-76 | 234.0 | L | Helix-turn-helix domain of resolvase | HTH_7,Resolvase |
| pAML2 | 2.18e-136 | 402.0 | K | DNA-binding transcription factor activity | MarR_2,ROK,RP-C,RP-C_C |
| pAML2 | 1.35e-267 | 751.0 | O | growth | - |
| pAML2 | 3.31e-82 | 249.0 | L | transposase activity | - |
| pAML2 | 1.32e-163 | 478.0 | S | Phage terminase large subunit (GpA) | Terminase_GpA |
| pAML2 | 1.6e-154 | 456.0 | S | Phage terminase | Terminase_GpA |

| | | | | | |
|---|---|---|---|---|---|
| | | | | large subunit (GpA) | |
| pAML2 | 1.8e-103 | 308.0 | - | - | - |
| pAML2 | 4.04e-32 | 114.0 | S | gpW | gpW |
| pAML2 | 3.06e-39 | 133.0 | S | Phage derived protein Gp49-like (DUF891) | Gp49 |
| pJPL24_01 | 0 | 60.5 | S | Plasmid stability protein | Arc |
| pJPL24_01 | 0 | 251 | S | Toxic component of a toxin-antitoxin (TA) module. An RNase | PIN |
| pJPL24_01 | 0 | 124 | K | Bacterial antitoxin of type II TA system, VapB | VapB_antitoxin |
| pJPL24_01 | 0 | 1747 | L | Conjugal transfer protein | AAA_30,MobA_MobL,Relaxase,Viral_helicase1 |
| pJPL24_01 | 0 | 205 | S | Conjugal transfer protein TraD | TraD |
| pJPL24_01 | 0 | 154 | S | PFAM Conjugal transfer TraD family protein | TraD |
| pJPL24_01 | 0 | 166 | S | CopG antitoxin of type II toxin-antitoxin system | CopG_antitoxin |
| pJPL24_01 | 0 | 136 | S | Ribonuclease toxin, BrnT, of type II toxin-antitoxin system | BrnT_toxin |
| pJPL24_01 | 0 | 646 | L | Plasmid replication initiator RepA | RPA |
| pJPL24_01 | 0 | 367 | D | involved in chromosome partitioning | AAA_31,CbiA |
| pJPL24_01 | 0 | 322 | L | Site-specific recombinases, DNA invertase Pin homologs | HTH_28,Resolvase |

| | | | | | |
|---|---|---|---|---|---|
| pJPL24_01 | 0 | 1253 | L | N-6 DNA Methylase | N6_Mtase |
| pJPL24_01 | 0 | 249 | V | Type I restriction modification DNA specificity domain | Methylase_S |
| pJPL24_01 | 0 | 115 | - | - | - |
| pJPL24_01 | 0 | 697 | L | Transposase | DDE_Tnp_1,DUF772 |
| pJPL24_01 | 0 | 393 | L | Site-specific recombinases, DNA invertase Pin homologs | Resolvase |
| pJPL24_01 | 0 | 419 | - | - | - |
| pJPL21 | 2.66e-284 | 794.0 | O | growth | - |
| pJPL21 | 4.71e-87 | 270.0 | L | Toprim domain | Toprim_3 |
| pJPL21 | 3.48e-11 | 60.5 | S | Putative antitoxin of bacterial toxin-antitoxin system, YdaS/YdaT | YdaS_antitoxin |
| pJPL21 | 1.8e-55 | 196.0 | D | Plasmid recombination enzyme | Mob_Pre |
| pJPL21 | 1.73e-281 | 775.0 | EGP | Major Facilitator Superfamily | MFS_1 |
| pJPL21 | 2.16e-69 | 219.0 | - | - | - |
| pJPL21 | 1.85e-66 | 209.0 | - | - | - |
| pJPL21 | 2.36e-116 | 337.0 | D | plasmid maintenance | AAA_31 |
| pJPL21 | 5.18e-51 | 166.0 | - | - | - |
| pJPL21 | 5.59e-23 | 93.6 | - | - | - |
| pJPL21 | 1.71e-11 | 73.2 | G | cellulose 1,4-beta-cellobiosidase activity | - |
| pJPL21 | 3.43e-70 | 219.0 | S | Tail protein | DUF2313 |
| pJPL21 | 2.16e-182 | 517.0 | S | Baseplate J-like protein | Baseplate_J |

| | | | | | |
|---|---|---|---|---|---|
| pJPL21 | 2.21e-61 | 194.0 | S | Phage protein GP46 | GP46 |
| pJPL21 | 1.04e-22 | 90.9 | K | Helix-turn-helix domain | HTH_3 |
| pJPL21 | 1.37e-79 | 240.0 | S | Bacteriophage Mu Gp45 protein | Phage_Mu_Gp45 |
| pJPL21 | 2.28e-171 | 489.0 | S | Mu-like prophage tail protein | Phage_GPD |
| pJPL21 | 7.63e-198 | 561.0 | S | DNA circularisation protein N-terminus | DNA_circ_N |
| pJPL21 | 4.06e-227 | 662.0 | S | phage tail tape measure protein | - |
| pJPL21 | 3.82e-82 | 243.0 | S | Phage tail tube protein | Tail_tube |
| pJPL21 | 2.29e-220 | 623.0 | S | Phage tail sheath protein subtilisin-like domain | Phage_sheath_1,Phage_sheath_1C |
| pJPL21 | 1.81e-30 | 115.0 | - | - | - |
| pJPL21 | 2.94e-178 | 508.0 | S | Phage major capsid protein E | Phage_cap_E |
| pJPL21 | 4.1e-54 | 173.0 | S | Bacteriophage lambda head decoration protein D | HDPD |
| pJPL21 | 2.44e-53 | 176.0 | OU | COG0616 Periplasmic serine proteases (ClpP class) | Peptidase_S49 |
| pJPL21 | 0.0 | 919.0 | S | COG5511 Bacteriophage capsid protein | Phage_portal_2 |
| pJPL21 | 8.82e-226 | 653.0 | S | Phage terminase large subunit (GpA) | Terminase_GpA |
| pJPL21 | 2.15e-39 | 134.0 | S | Phage derived protein Gp49-like (DUF891) | Gp49 |
| pJPL21 | 3.79e-44 | 145.0 | S | gpW | gpW |

| | | | | | |
|---|---|---|---|---|---|
| pJPL21 | 1.01e-133 | 395.0 | K | DNA-binding transcription factor activity | MarR_2,ROK,RP-C,RP-C_C |
| pJPL21 | 2.18e-75 | 231.0 | L | Helix-turn-helix domain of resolvase | HTH_7,Resolvase |
| pJPL21 | 6.48e-21 | 89.7 | K | Transcriptional regulator | RepL |
| pJPL24_02 | 1.15e-94 | 285.0 | L | Initiator Replication protein | Rep_3 |
| pJPL24_02 | 8.65e-38 | 129.0 | K | copG family | RHH_1 |
| pJPL24_02 | 6.93e-41 | 136.0 | S | Plasmid stabilization | ParE_toxin |
